# Comparison of Tug-of-War Models Assuming Moran versus Branching Process Population Dynamics

**DOI:** 10.1101/2023.10.20.563302

**Authors:** Khanh N. Dinh, Monika K. Kurpas, Marek Kimmel

## Abstract

Mutations arising during cancer evolution are typically categorized as either ‘drivers’ or ‘passengers’, depending on whether they increase the cell fitness. Recently, McFarland et al. introduced the Tug-of-War model for the joint effect of rare advantageous drivers and frequent but deleterious passengers. We examine this model under two common but distinct frameworks, the Moran model and the branching process. We show that frequently used statistics are similar between a version of the Moran model and the branching process conditioned on the final cell count, under different selection scenarios. We infer the selection coefficients for three breast cancer samples, resulting in good fits of the shape of their Site Frequency Spectra. All fitted values for the selective disadvantage of passenger mutations are nonzero, supporting the view that they exert deleterious selection during tumorigenesis that driver mutations must compensate.

## 1 Introduction

As demonstrated in the seminal paper [12], mutations in different cancers vary substantially in counts and patterns. These differences reflect distinct defects in DNA repair mechanisms, cancer exposures, and cell types. The authors also reported evidence for ‘driver’ mutations in about 120 genes, which contribute to tumorigenesis. However, the majority of somatic mutations are likely ‘passengers’, which do not have an effect on tumor progression. In the language of population genetics, driver mutations are selectively advantageous to cancers, while the passengers are at best neutral.

The concept of a model involving the joint effect of rare advantageous and frequent neutral or slightly deleterious mutations can be applied to describe evolution of cancer genes. To the best of our knowledge, such model was first introduced by McFarland and co-authors, in a series of publications [18, 19, 20], and named the Tug-of-War model to reflect the competition between driver and passenger mutations.

The original Tug-of-War model [18] assumed that the cell death rate increases with the number of cells in the population increasing, which creates a mechanism for limiting the eventual tumor size. In other papers [16, 17], a Moran model was used for the population process, which provides a strict bound on the number of proliferating cells (see relevant discussion in [16]). Another assumption of [18] was that driver mutations become instantly fixed in the population, which may be acceptable under very strong selection (for mathematical details, see Bobrowski et al. [2]), but in general it is not satisfied.

The literature includes many examples of comparisons of how mutation, drift and selection interact in different population dynamics frameworks such as branching process versus Moran model [3, 4] or Wright-Fisher model with population of varying size [5, 6]. Two possible modes of selection in cell populations are “crowding out” in which a faster-growing clone makes the slower-growing one rare to the point of being negligible, or “competitive replacement”, in which individual cells inhibit each other’s replacement by a descendant. Supercritical branching process models lead to the former, while the Moran and Wright-Fisher models to the latter. A version interpolating between these two mechanisms is the well-cited Gerrish and Lenski model [11]. We will return to these models in the Discussion.

In the present paper, we compare the Tug-of-War in the multitype Moran model with constant population size and a critical multitype branching process. The latter is conditional on non-extinction or other restrictions. We explore similarities and differences between the two types of selection in cell populations. This contributes to the ongoing discussion of which models are most appropriate for proliferating cell populations under drift, mutation, and selection.

We begin with mathematical definitions of the two versions of Tug-of-War process. Then we present simulation results, which demonstrate the differences between the long-term behavior of the two versions under different selection scenarios. Finally, we infer the selection coefficients for some breast cancer samples using the Moran framework, and cross-examine the fitted parameters with the branching process.

## 2 Models and Data

### 2.1 Moran process Tug-of-War models

Model descriptions in this section, are essentially summaries of the descriptions in [16] and to avoid redundancy, we summarize only the most essential features. For more detailed descriptions, see [16].

#### 2.1.1 Model A

In this model (**Figure 1A**), we contextualize the Tug-of-War within the Moran model framework with multiple allelic types. We examine a population with a constant size of *N* cells. Each cell *i* is described by a pair of integers *γ*_*i*_ = (*α*_*i*_, *β*_*i*_), where *α*_*i*_ and *β*_*i*_ represent the quantities of drivers and passengers in its genotype, respectively. The cell’s fitness is then defined as

**Figure 1:**
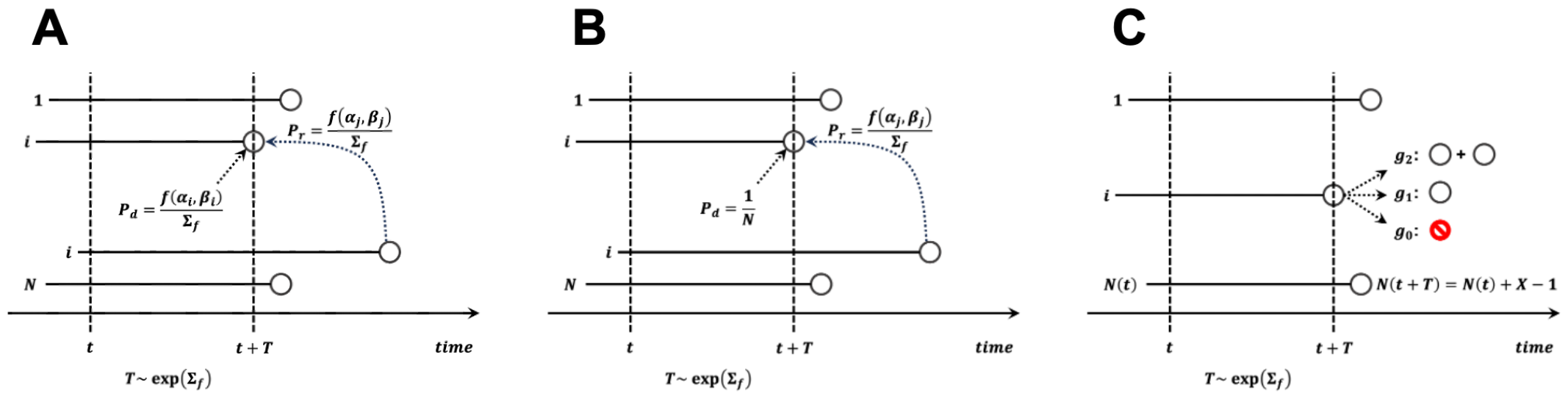
Graphical depiction of cell death and division events in **(A)** Moran A model, **(B)** Moran B model, and **(C)** branching process. **Notation for all models:** *t*, current time; *T*, time to next event; *f* (*α, β*), fitness of cell with *α* drivers and *β* passengers; Σ_*f*_ =Σ_*k*_ *f* (*α*_*k*_, *β*_*k*_). **Notation for Moran models:** *N*, constant cell count in the process; *i*, cell dying and to be replaced; *j*, cell replacing cell *i*; **Notation for branching process:** *N* (*t*), cell count at time *t*; *i*, cell chosen for division; {*g*_0_, *g*_1_, *g*_2_}, progeny count distribution; *X*, progeny count of cell *i*; *N* (*t* + *T*), cell count at time *t* + *T* after division event.

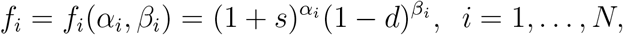

where *s >* 0 is the selective advantage associated with a driver mutation, while *d ∈* (0, 1) represents the selective disadvantage of a passenger mutation (selection coefficients, of driver and passenger mutations; see the Natural Selection chapter of the book by Durrett [9]). Under the time-continuous Markov Chain assumption, the time until the next death - replacement event is exponentially distributed with parameter Σ_*f*_ =Σ _*i∈{*1,…,*N }*_ *f*_*i*_. The dying cell *i* is selected from a distribution biased by fitness, i.e., with probability mass function (pmf) {*f*_*i*_*/*Σ_*f*_, *i* = 1, …, *N }*. The cell *j* that replaces the dying cell is selected from the same distribution.

The time until the next mutation event is exponentially distributed with parameter N*μ*, where *μ* is the mutation rate per cell. The cell selected to mutate is chosen uniformly among the *N* cells. Its state then changes from (α, *β*) to (α + 1, *β*) with probability *p ∈* (0, 1), or to (α, *β* + 1) with probability 1 *− p*. In combination, the time to the next event is random and exponentially distributed with parameter

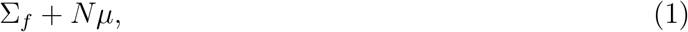

called the total rate of death - replacement and mutation events.

#### 2.1.2 Model B

Model B (**Figure 1B**) is defined similarly to Model A, with the time until the next death - replacement event being exponentially distributed with parameter Σ_*f*_. However, instead of being biased by fitness, the dying cell is chosen among all *N* cells uniformly. The time until the next event is likewise exponentially distributed with the same parameter in Equ. (1).

#### 2.1.3 Model A versus Model B

As we noted in [16], the crucial difference between Model A and Model B lies in the expected fitness increment in the population after the death - replacement event. This is equal to the difference of *f*_*j*_ *− f*_*i*_, where *f*_*i*_ and *f*_*j*_ are the fitnesses of the dead cell and the replacing cell, in the absence of mutations. The mutation-selection balance condition was derived in [2] in the form of

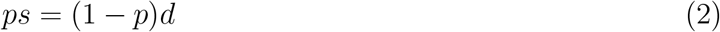

When *ps >* (1 *− p*)*d*, drivers “dominate” over passengers, and the reverse occurs if *ps <* (1 *− p*)*d*. The expected fitness is intact after the death - replacement process in Model A, hence the expected fitness trend follows the mutation-selection condition. In Model B, the outcome is more complex (see Results and Discussion).

### 2.2 Branching process model

For the branching process model (**Figure 1C**), we consider a population consisting of *N* (*t*) cells at time *t*. Similar to the Moran models, the fitness of a cell *i* of type (*α*_*i*_, *β*_*i*_) with *α*_*i*_ drivers and *β*_*i*_ passengers is defined by *f*_*i*_ = (1 + *s*) ^*αi*^ (1 *− d*) ^*βi*^.

Two possible event types can occur in the branching process: cell division and mutation. The time to the next cell division is exponentially distributed with rate Σ_*f*_ = Σ_*i∈{*1,…,*N* (*t*)}_ *f*_*i*_, fitness sum of all *N* (*t*) cells. The dividing cell is chosen from the *N* (*t*) cells with probability weighted by fitness {*f*_*i*_*/*Σ_*f*_ }, and its progeny count follows a given pmf {*g*_0_, *g*_1_, … }. If the progeny count is 0, the population loses the chosen cell. On the other hand, it survives if the progeny count is 1, and multiplies if the progeny count is more than 1. Additionally, the time to the next mutation event is exponentially distributed with rate *μN* (*t*). A uniformly chosen cell acquires a driver and changes its type to (*α* + 1, *β*) with probability *p*, or it acquires another passenger to become type (*α, β* + 1) with probability 1 *− p*.

#### 2.2.1 Criticality and conditioning

One important difference between the Moran and branching process settings is that the total cell count at any time remains constant at *N* in the Moran models. The branching process model starts from the same cell count, i.e. *N* (0) = *N*, but it can change at random throughout time. For a direct comparison to Moran models, we may assume that 𝔼 (*N* (*t*)) = *N*. This is satisfied if we require that the mean progeny count of any cell is 1, i.e. that the branching process is critical [1, 15].

Even with criticality, at any time *N* (*t*) can fixate at 0 (in which case the process enters extinction) or increase to a larger count. The probability of extinction increases as either *t* increases ([15], Section 3.3) or the cell fitness increases as a result of time scale change. We analyze the effects of two types of conditioning on the results of the branching process. The first type conditions the branching process on non-extinction, meaning *N* (*t*_*f*_) *>* 0 at the final time *t*_*f*_. The second type imposes a more stringent condition on the branching process, requiring that *N* (*t*_*f*_) *∈* [*N − c, N* + *c*] for some small constant *c*.

#### 2.2.2 Distribution of progeny cell counts

We also study the impact of the distribution of progeny cell count {*g*_*k*_, *k* = 0, 1, … } on the outcomes of the branching process. In this study, we impose *g*_*k*_ = 0 for *k >* 2. Therefore, a chosen cell dies if *k* = 0, remains unchanged if *k* = 1, or divides into two progeny cells if *k* = 2. The criticality requirement, discussed in the previous section, is satisfied if *g*_0_ = *g*_2_. Note that the branching process setting discussed in this paper is equivalent to a birth-death process where each cell *i* with fitness *f*_*i*_ dies with rate *g*_0_*f*_*i*_ or divides with rate *g*_2_*f*_*i*_.

A common progeny count distribution is such that *g*_0_ = *g*_2_ = 0.25 and *g*_1_ = 0.5, which is equivalent to a binomial distribution with rates *n* = 2, *p* = 0.5. For a direct comparison in simulations to the Moran models, the fitness in the branching process is scaled up by 4. Hence, the wait time until the next division event of a cell with fitness *f*_*i*_ is exponentially distributed with rate 4*f*_*i*_. This event has equal probability to be a cell death or a cell division, both at 0.25. Therefore, the model is equivalent to a birth-death process, where the birth and death rates are 0.25 *×* 4*f*_*i*_ = *f*_*i*_. In comparison, in the Moran models, the death - replacement events also occur at rate *f*_*i*_, resulting in two cells being chosen to divide and die, respectively. Because the event rates are now similar, it is easier to directly compare the model behaviors under different selection scenarios.

We will also investigate the effect of changing the progeny cell count distribution {*g*_0_, *g*_1_, *g*_2_}. First, we retain the criticality by assuming *g*_0_ = *g*_2_, and analyze the branching process with different values for *g*_1_. We set *g*_0_ = *g*_2_ = 0.5 and *g*_1_ = 0, which doubles the probabilities for cell division and death events. This is similar to increasing the birth and death rates in a birth-death process, hence we name this model “fast BP”. We also consider *g*_0_ = *g*_2_ = 0.05 and *g*_1_ = 0.9 (“slow BP”), which decreases the cell division and death probabilities. Second, we investigate the supercritical branching process by setting *g*_2_ *> g*_0_. For each of these parameter sets, we will analyze the differences in sample statistics, compared to the binomial branching process and the Moran models.

### 2.3 Site Frequency Spectrum

As in other preceding paragraphs, in this section, we include a summary of the descriptions in [16]. For more details, see [16].

One of the common summary statistics of the sequence data is the so-called Site Frequency Spectrum. In a sequencing experiment with *n* cells, we can estimate for each novel somatic mutation call the number of cells carrying that mutation. The number *S*_*n*_(*k*) of mutations present in *k* cells is put into a vector (*S*_*n*_(1), *S*_*n*_(2), …, *S*_*n*_(*n −* 1)) called the Site Frequency Spectrum, abbreviated to SFS. Frequently number of cells that were sampled is not know, as for example in the bulk sequencing data. However, we can estimate the relative proportion of the mutant at each site, and so arrive at a frequency spectrum based on proportions with notation *S*(*x*) = *S*(*k/n*), where *x* is treated as a continuous variable, such that *x ∈* (0, 1) (or *x ∈* (0, 1*/*2) if we consider diploid genome). The SFSs *S*(*k*) and *S*(*x*) are idealized versions of the empirical variant allele frequency (VAF) graph. It is convenient for reasons explained in [16] (Section 2.4) to employ the cumulative tail of the SFS *S*(*x*)

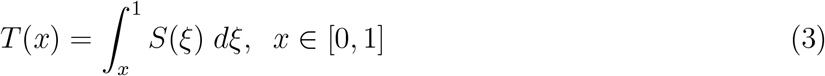

### 2.4 DNA Sequencing of Cell Samples from Breast Cancer Specimens

#### 2.4.1 DNA sample collection and processing

Most of the details of DNA sample collection and processing in this section, overlap with those in [16] and we refer to the description there. We only mention here several basic details. Tissue samples from primary breast tumor were collected at the Department of Applied Radiology of the Maria Sklodowska-Curie National Research Institute of Oncology, Krakow Branch in Poland. The set of tumor and normal control samples called specimen G2 is HER2+ breast cancer, while sets described as G32 and G41 are triple-negative breast cancer type and luminal A type, respectively.

#### 2.4.2 Removal of FFPE artifacts

Fixation of tissues in formalin leads to deamination of cytosine to uracil, which can be recognized by sequencing as C*>*T or G*>*A type modifications [8].

A significant portion of the variants detected in our WES data are a possible artifact of sample fixation in formalin. This is indicated primarily by the statistics of the number of variants of a specific type, where C:G*>*T:A definitely dominates (Table 1).

**Table 1:** Statistics of the number of variants of a specific type.

| Patient ID | C:G>A:T | C:G>G:C | C:G>T:A | T:A>A:T | T:A>C:G | T:A>G:C | sum |
| --- | --- | --- | --- | --- | --- | --- | --- |
| G2 | 190 | 78 | 8379 | 87 | 430 | 84 | 9248 |
| G32 | 86 | 47 | 6394 | 56 | 195 | 36 | 6814 |
| G41 | 53 | 42 | 2088 | 44 | 176 | 38 | 2441 |

The reason for such a large number of this type of changes may be the duplication of variants related to deamination in PCR amplification (necessary in the case of WES, especially in the case of samples with a low amount of DNA).

Omitting all C:G*>*T:A variants would result in the loss of approximately 1/6 of the true variants. However, information about the frequency of reads with a specific orientation can be used to identify variants associated with the method of sample fixation. For this purpose, the SOBDetector [7] program was used. The software is based on the fact that formalin fixation most likely affects only one of the DNA strands (the C:G pair becomes the T:G pair) and therefore the paired-end next-generation sequencing approach can help this additional filtering step. By counting not only the number of reads supporting alternative alleles, but also the relative orientation of the reads (Forward-Reverse:FR or Reverse-Forward:RF), these FFPE artifacts will likely have a strand orientation bias toward one of the directions, while true mutations should have approximately the same number of FR and RF reads.

This work uses data in which the expected FFPE artifacts have been filtered out by SOBDetector.

## 3 Results

### 3.1 Behavior of Moran and branching process models in extreme cases

We investigate the similarities and differences between Moran and branching process models. One thousand simulations are performed for each model, and each simulation starts with *N* = 100 cells with no mutations at *t*_0_ = 0 under different values for the selection coefficients *s* and *d*, mutation rate *μ* and probability *p* of driver mutations (or equivalently probability 1 *− p* of passenger mutations). We then examine the statistics at final time *t*_*f*_ = 100 as well as during the entire time line [*t*_0_, *t*_*f*_].

Five versions of branching processes are studied. This includes the branching process with *g*_0_ = *g*_2_ = 0.25 and *g*_1_ = 0.5 conditioned on non-extinction (yellow), and the same branching process conditioned on the final population *N* (*t*_*f*_) restricted in [90, 110] (orange). These models are termed “binomial BP” in the comparisons, since the progeny cell count distribution is binomial in this case. We also include the branching process with *g*_0_ = *g*_2_ = 0.5, *g*_1_ = 0 (purple, termed “fast BP”) and *g*_0_ = *g*_2_ = 0.05, *g*_1_ = 0.9 (cyan, termed “slow BP”), both conditioned on *N* (*t*_*f*_) *∈* [90, 110].

The fifth branching process model is conditioned on non-extinction with *g*_0_, *g*_1_ and *g*_2_ computed such that the population size is expected to double between [*t*_0_, *t*_*f*_] under neutral evolution. It can be shown that *g*_0_ and *g*_2_ are required to satisfy

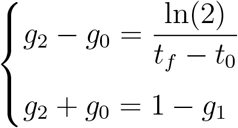

We choose *g*_1_ = 0.5, resulting in *g*_0_ = 0.2465 and *g*_2_ = 0.2535. This model is referred to as “supercritical BP” (gray) in the numerical comparisons. Finally, Moran A and Moran B are represented in dark blue and green, respectively.

Since we scale up the fitness by 4 in the branching process models to make them similar to the Moran models, in the results we scale the fitnesses down by 4, for more convenient comparisons. The division count, defined to be the total number of cell division events observed in a simulation, increases linearly with the total cell count and cell fitness, i.e. the rate at which cells divide. Since the expected cell count in a critical BP is identical to the Moran models, the division count in BP is 4 times higher than in the Moran models due to the fitness being scaled up. Therefore, we also downscale the BP division count by 4 to directly compare between different models.

#### 3.1.1 Neutral evolution case

**Figure 3** presents the simulated results for *s* = *d* = 0, which implies that all mutations are neutral, *μ* = 0.1 and *p* = 1*/*11.

**Figure 2:**
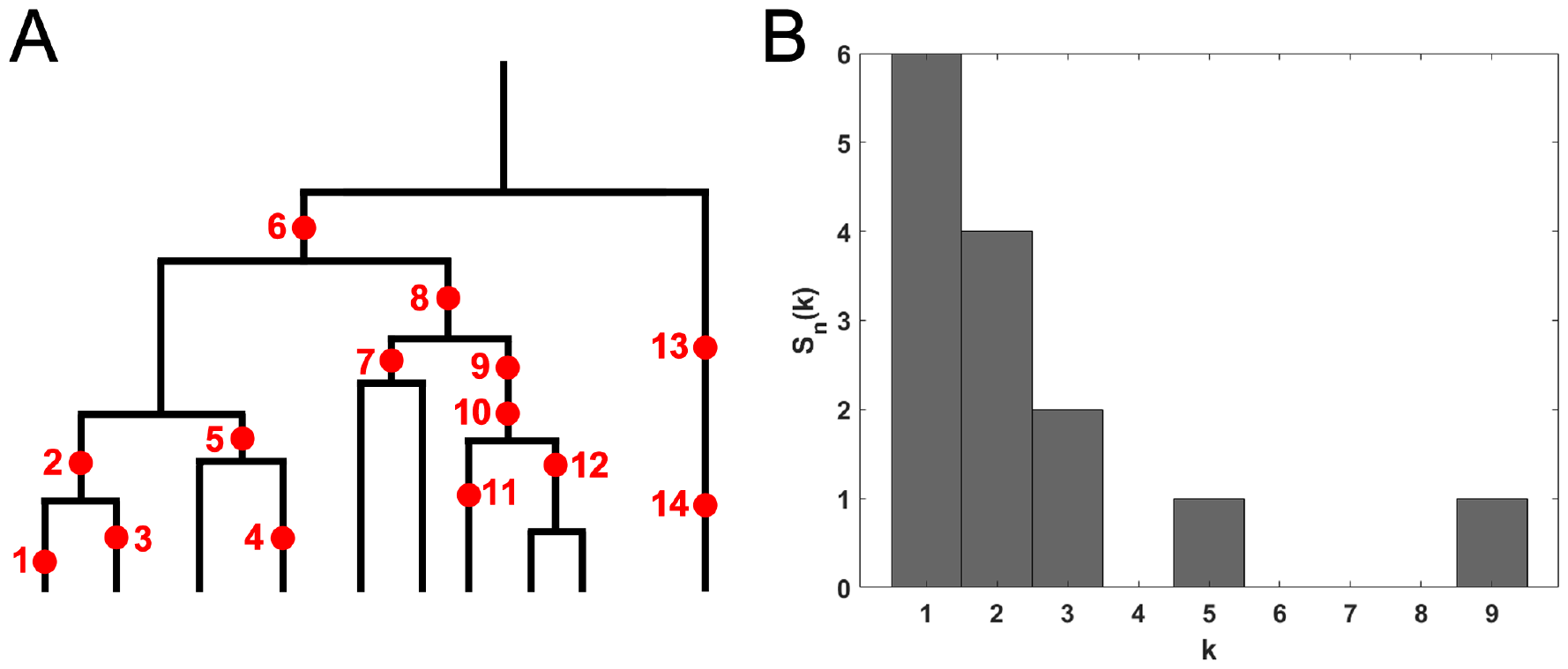
The Site Frequency Spectrum (SFS). **(A)** Genealogy of a sample of *n* = 10 cells includes 14 mutational events, denoted by red dots. Time is running down the page. Mutations 1, 3, 4, 11, 13, and 14 (total of 6 mutations) are present in a single cell, mutations 2, 5, 7 and 12 (total of 4 mutations) are present in two cells, mutations 9 and 10 (2 mutations) are present in three cells, mutation 8 (1 mutation) is present in five cells and mutation 6 (1 mutation) is present in 9 cells. **(B)** The resulting site frequency spectrum, *S*_10_(1) = 6, *S*_10_(2) = 4, *S*_10_(3) = 2, *S*_10_(5) = 1, and *S*_10_(9) = 1, other *S*_*n*_(*k*) equal to 0.

**Figure 3:**
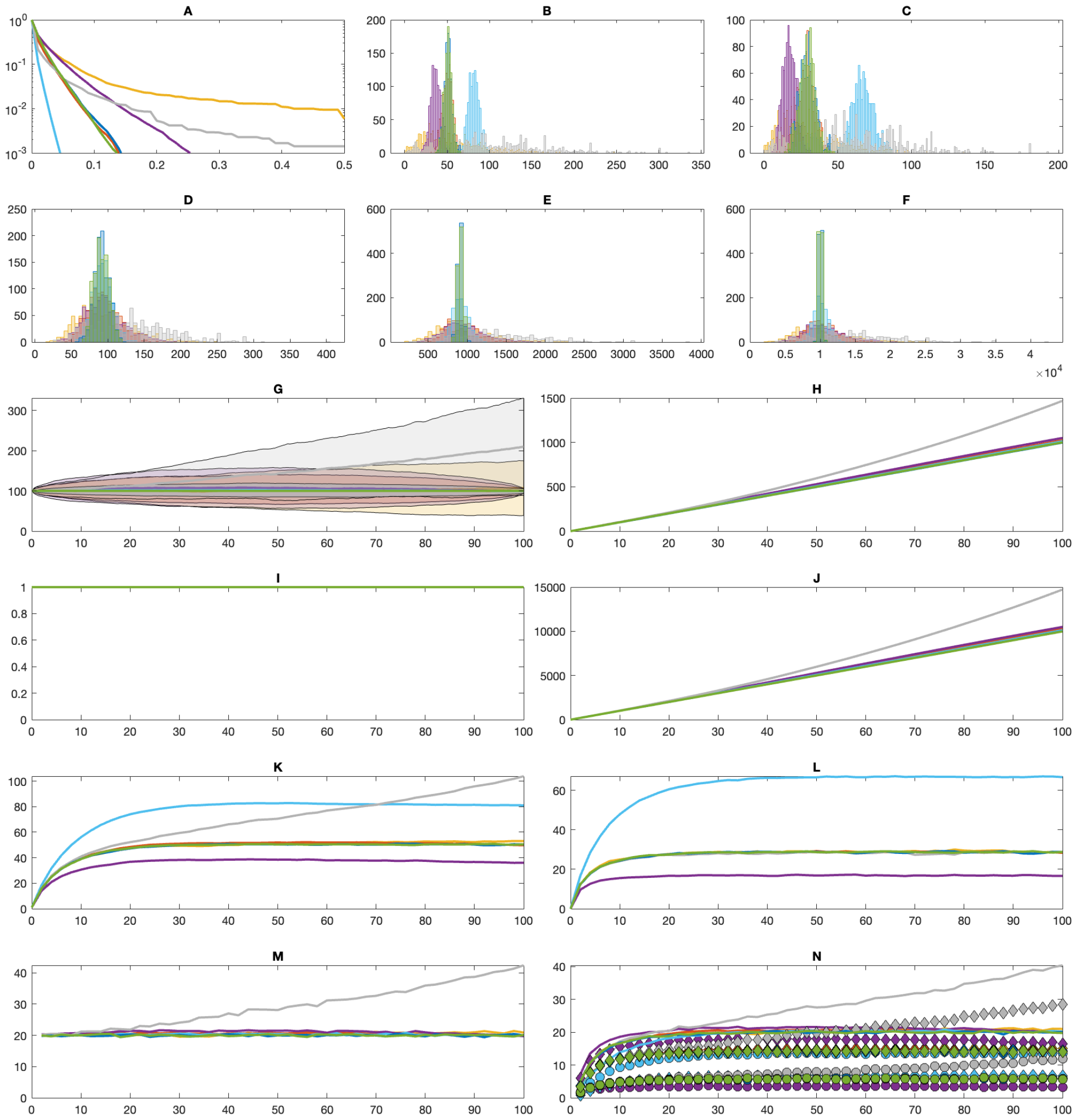
Comparisons between Moran and branching process (BP) models in the “neutral” setting. **(A)** Average cumulative tail of the mutational Site Frequency Spectra. **(B)** Distributions of allele counts at *t*_*f*_. **(C)** Distributions of singleton counts at *t*_*f*_. **(D-F)** Distributions of counts of driver mutations **(D)**, passenger mutations **(E)** and divisions **(F)** within [*t*_0_, *t*_*f*_]. **(G-N)** Trajectories of the averages over time of population sizes (+/-std) **(G)**, cumulative mutation counts **(H)**, fitness **(I)**, cumulative division/replacement counts **(J)**, allele counts **(K)**, percentage of singletons among all alleles **(L)**, allele birth counts **(M)** and allele death counts **(N)**. Allele death counts (lines) are categorized into mutation events (circles) and division/replacement events (diamonds). Dark blue = Moran A, green = Moran B, yellow = “binomial BP” with *g*_0_ = *g*_2_ = 0.25 (non-extinction), orange = “binomial BP” with *g*_0_ = *g*_2_ = 0.25 (*N* (*t*_*f*_) *∈* [90, 110]), purple = “fast BP” with *g*_0_ = *g*_2_ = 0.5 (*N* (*t*_*f*_) *∈* [90, 110]), cyan = “slow BP” with *g*_0_ = *g*_2_ = 0.05 (*N* (*t*_*f*_) *∈* [90, 110]), gray = “supercritical BP” with *g*_0_ = 0.2465, *g*_1_ = 0.5 and *g*_2_ = 0.2535 (non-extinction).

Since all cells have the same fitness, the formulations for Moran A and Moran B are identical. This is reflected in identical distributions among all statistics among the two variations (see Appendix C, Table 2, dark blue for Moran A and green for Moran B). Moreover, the binomial BP conditioned on *N* (*t*_*f*_) *∈* [90, 110] also has the same distributions of allele counts and singleton counts, albeit with slightly higher variances (at final time in **Figure 3B-C** and throughout history in **Figure 3K-L**). The higher variances originate from wider distributions of event counts in the BP compared to Moran, these latter stemming from the fact that total cell count in BP varies throughout time, as opposed to Moran where the total cell count remains constant (**Figure 3D-G**). Note that even though the statistics have higher variances, their averages are similar to the Moran models both throughout time and at the final time (Appendix C, Table 2, orange).

**Table 2:** Statistics (mean *±* standard deviation) for Figure 3 (neutral evolution). **Model color code:** dark blue = Moran A, green = Moran B, yellow = “binomial BP” with *g*_0_ = *g*_2_ = 0.25 (nonextinction), orange = “binomial BP” with *g*_0_ = *g*_2_ = 0.25 (*N* (*t*_*f*_) *∈* [90, 110]), purple = “fast BP” with *g*_0_ = *g*_2_ = 0.5 (*N* (*t*_*f*_) *∈* [90, 110]), cyan = “slow BP” with *g*_0_ = *g*_2_ = 0.05 (*N* (*t*_*f*_) *∈* [90, 110]), gray = “supercritical BP” with *g*_0_ = 0.2465, *g*_1_ = 0.5 and *g*_2_ = 0.2535 (non-extinction). **Statistics:** B = allele counts at *t*_*f*_, C = singleton counts at *t*_*f*_, D = driver mutation count within [*t*_0_, *t*_*f*_], E = passenger mutation count within [*t*_0_, *t*_*f*_], F = division count within [*t*_0_, *t*_*f*_].

| Model | B | C | D | E | F |
| --- | --- | --- | --- | --- | --- |
| Dark blue | 50.3±4.6 | 28.7±5.36 | 91.1±9.9 | 907.6±31.3 | 10005.5±207.0 |
| Green | 50.4±4.5 | 28.7±5.0 | 90.8±9.8 | 909.9±31.4 | 9992.3±206.1 |
| Yellow | 53.0±33.6 | 29.8±19.5 | 93.3±36.0 | 930±355.9 | 10235.7±3888.0 |
| Orange | 49.7±6.1 | 28.2±5.8 | 93.5±21.5 | 929.8±194.1 | 10261.6±2121.3 |
| Purple | 36.0±5.9 | 16.6±4.5 | 95.3±27.9 | 954.8±261.3 | 10498.6±2862.7 |
| Cyan | 81.0±6.2 | 66.5±7.5 | 92.1±13.3 | 919.9±95.0 | 10117.1±1003.4 |
| Gray | 103.8±58.7 | 58.9±33.8 | 133.1±53.4 | 1335.8±524.3 | 14723.8±5718.5 |

The impact of relaxing the conditioning of BP can be observed by comparing binomial BP conditioned on *N* (*t*_*f*_) *∈* [90, 110], binomial BP conditioned on non-extinction and supercritical binomial BP with *g*_0_ = 0.2465, *g*_1_ = 0.5 and *g*_2_ = 0.2535, conditioned on non-extinction. As the population size can vary widely throughout time if conditioned only on non-extinction (**Figure 3G**), every statistics in the comparison has much higher variances (Appendix C, Table 2, yellow for critical BP and gray for supercritical BP). However, for critical BP, the averages remain faithful to both Moran models and the more stringently conditioned BP. Only in the case of supercritical BP do all statistics differ, even cumulative mutation count (**Figure 3H**) and cumulative division/replacement count (**Figure 3J**), which is strictly associated with rapidly growing population size.

We then evaluate the impact of changing the progeny cell count distributions, while retaining criticality. The fast BP has increased *g*_0_ and *g*_2_ and therefore is equal to a birth-death process with higher rates. This leads to higher variances in the population size throughout time compared to the binomial BP, even if similarly conditioned (**Figure 3G**, Appendix C: Table 2, purple). Importantly, the fast BP also results in both less alleles and lower percentage of singletons within all alleles (**Figure 3B-C, K-L**). Conversely, the slow BP is equivalent to a birth-death process with lower rates, whose population size therefore varies less than the binomial BP (**Figure 3G**). Both its allele count and percentage of singleton count is much higher than in the binomial BP (**Figure 3B-C, K-L**, Appendix C: Table 2, cyan).

Figure 3N details the rates at which alleles are lost from the population, divided into two categories: (a) cell deaths (division with no progeny cells in BP, or replacement in Moran), and (b) cell mutation (where the only remaining cell that carries the allele mutates into a different allele). The allele death counts, either combined or categorized to (a) or (b), are similar in all cases except supercritical BP and fast or slow BP. The rate of allele death is slightly lower for slow BP and slightly higher in the case of fast BP. They also have different categorized allele death counts. Compared to binomial BP, alleles are removed more frequently due to mutations in slow BP. Conversely, they are more likely to be removed due to failed divisions in fast BP. The supercritical BP is the only case in which allele death count does not reach plateau and is still increasing during the simulated time period, as the population grows exponentially.

#### 3.1.2 Balanced evolution

**Figure 4** showcases the simulated results for *s* = 0.1, *d* = 0.01, *μ* = 0.1 and *p* = 1*/*11. As *ps* = (1 *− p*)*d*, condition (2) is satisfied, therefore the average fitness remains constant for Moran A model (**Figure 4I**). Moreover, the fitness in binomial BP also remains unchanged on average over time. As a result, the distributions for all statistics are similar between the balanced evolution setting and the neutral evolution setting (Appendix B, Table 3), discussed in the last section, for Moran A and binomial BP. The outcomes following modulation of the progeny cell count distribution or relaxing the conditioning for BP, likewise remain unchanged compared to the neutral evolution setting.

**Table 3:**
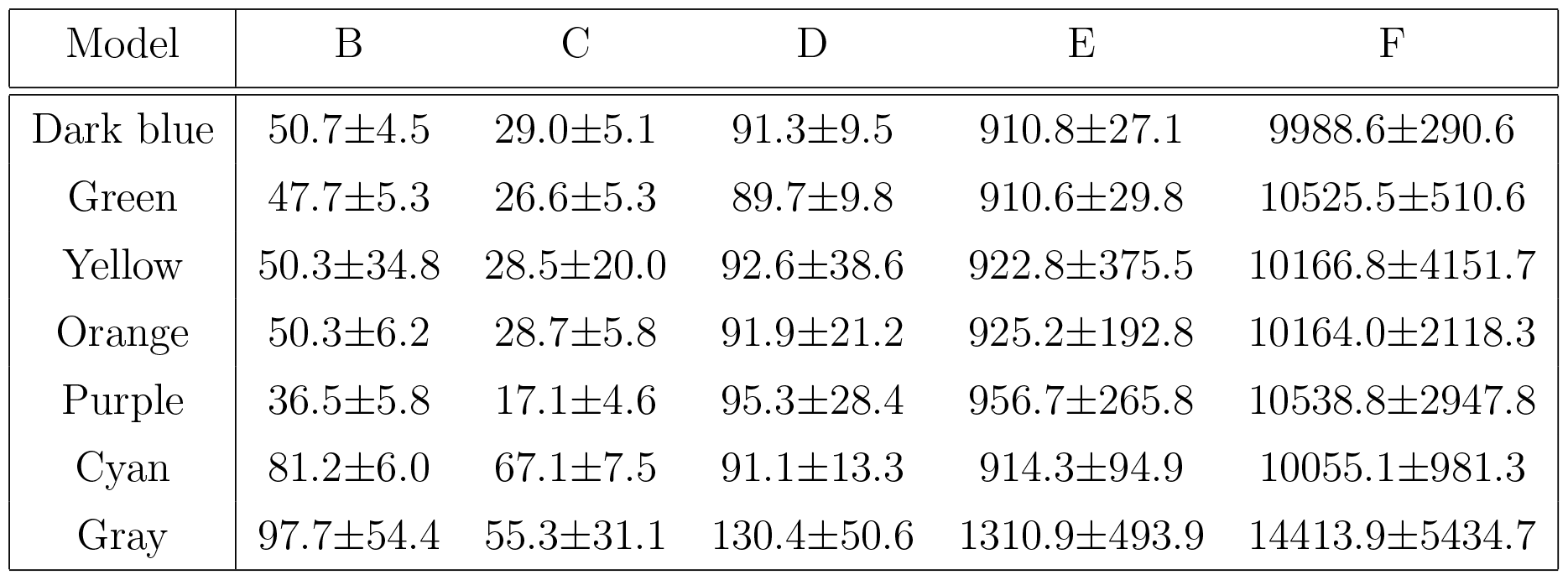
Statistics for Figure 4 (balanced evolution)

| Model | B | C | D | E | F |
| --- | --- | --- | --- | --- | --- |
| Dark blue | 50.7±4.5 | 29.0±5.1 | 91.3±9.5 | 910.8±27.1 | 9988.6±290.6 |
| Green | 47.7±5.3 | 26.6±5.3 | 89.7±9.8 | 910.6±29.8 | 10525.5±510.6 |
| Yellow | 50.3±34.8 | 28.5±20.0 | 92.6±38.6 | 922.8±375.5 | 10166.8±4151.7 |
| Orange | 50.3±6.2 | 28.7±5.8 | 91.9±21.2 | 925.2±192.8 | 10164.0±2118.3 |
| Purple | 36.5±5.8 | 17.1±4.6 | 95.3±28.4 | 956.7±265.8 | 10538.8±2947.8 |
| Cyan | 81.2±6.0 | 67.1±7.5 | 91.1±13.3 | 914.3±94.9 | 10055.1±981.3 |
| Gray | 97.7±54.4 | 55.3±31.1 | 130.4±50.6 | 1310.9±493.9 | 14413.9±5434.7 |

**Figure 4:**
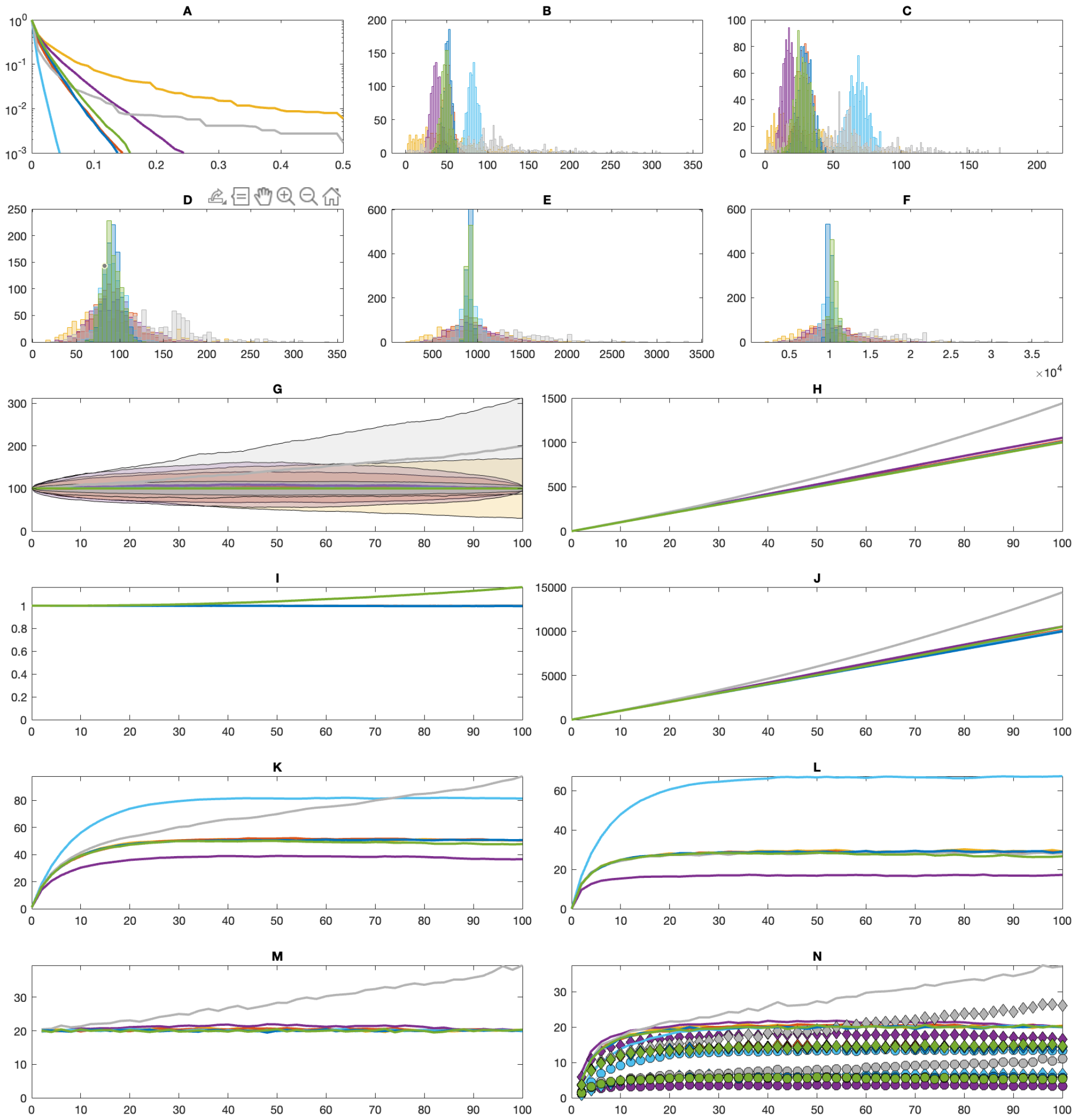
Comparisons between Moran and branching process (BP) models in the “balanced” setting. **(A)** Average cumulative tail of the mutational Site Frequency Spectra. **(B)** Distributions of allele counts at *t*_*f*_. **(C)** Distributions of singleton counts at *t*_*f*_. **(D-F)** Distributions of counts of driver mutations **(D)**, passenger mutations **(E)** and divisions **(F)** within [*t*_0_, *t*_*f*_]. **(G-N)** Trajectories of the averages over time of population sizes (+/-std) **(G)**, cumulative mutation counts **(H)**, fitness **(I)**, cumulative division/replacement counts **(J)**, allele counts **(K)**, percentage of singletons among all alleles **(L)**, allele birth counts **(M)** and allele death counts **(N)**. Allele death counts (lines) are categorized into mutation events (circles) and division/replacement events (diamonds). Dark blue = Moran A, green = Moran B, yellow = “binomial BP” with *g*_0_ = *g*_2_ = 0.25 (non-extinction), orange = “binomial BP” with *g*_0_ = *g*_2_ = 0.25 (*N* (*t*_*f*_) *∈* [90, 110]), purple = “fast BP” with *g*_0_ = *g*_2_ = 0.5 (*N* (*t*_*f*_) *∈* [90, 110]), cyan = “slow BP” with *g*_0_ = *g*_2_ = 0.05 (*N* (*t*_*f*_) *∈* [90, 110]), gray = “supercritical BP” with *g*_0_ = 0.2465, *g*_1_ = 0.5 and *g*_2_ = 0.2535 (non-extinction).

However, the fitness in Moran B increases over time instead of remaining constant (**Figure 4I**), which leads to slightly higher division count (**Figure 4F**) even though the mutation count, depending only on the population size which remains constant, is unchanged (**Figure 4D-E**). This results in slightly lower allele count and singleton count (**Figure 4B-C**), indicative of selective pressure.

#### 3.1.3 Driver domination case

To understand the consequences when the driver mutations are strongly selectively advantageous, we set *s* = 0.25, *d* = 0, *μ* = 0.1 and *p* = 1*/*10 and compare the statistics in **Figure 5** and in Appendix B, Table 4.

**Table 4:** Statistics for Figure 5 (driver domination)

| Model | B | C | D | E | F |
| --- | --- | --- | --- | --- | --- |
| Dark blue | 45.4±5.2 | 24.2±5.2 | 100.1±9.8 | 899.7±30.6 | 11455.7±772.4 |
| Green | 23.6±7.8 | 9.4±4.8 | 100.4±9.7 | 899.2±28.3 | 20767.0±7693.0 |
| Yellow | 44.6±29.5 | 24.0±16.2 | 98.2±37.9 | 888.3±331.5 | 11153.3±4416.3 |
| Orange | 45.3±6.3 | 24.3±5.5 | 103.2±25.2 | 928.1±207.0 | 11683.1±2716.7 |
| Purple | 32.3±5.9 | 14.1±4.5 | 105.0±33.3 | 942.4±287.8 | 11818.8±3751.9 |
| Cyan | 77.8±6.0 | 61.7±7.2 | 100.5±14.2 | 903.9±94.0 | 11441.5±1242.2 |
| Gray | 97.3±56.8 | 51.6±29.8 | 150.2±58.5 | 1341.1±506.0 | 17454.9±7228.5 |

**Figure 5:**
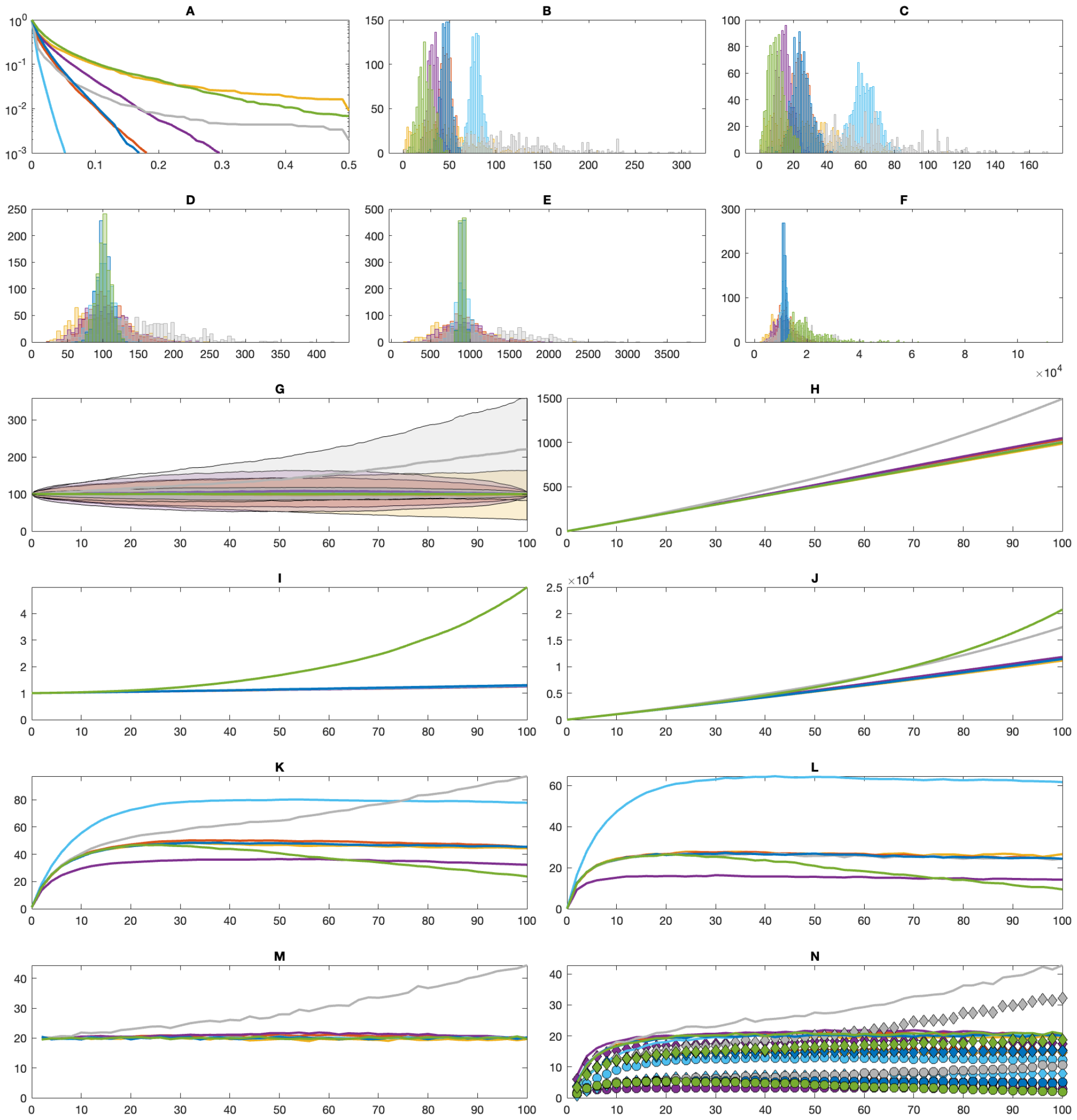
Comparisons between Moran and branching process (BP) models in the “selective” setting. **(A)** Average cumulative tail of the mutational Site Frequency Spectra. **(B)** Distributions of allele counts at *t*_*f*_. **(C)** Distributions of singleton counts at *t*_*f*_. **(D-F)** Distributions of counts of driver mutations **(D)**, passenger mutations **(E)** and divisions **(F)** within [*t*_0_, *t*_*f*_]. **(G-N)** Trajectories of the averages over time of population sizes (+/-std) **(G)**, cumulative mutation counts **(H)**, fitness **(I)**, cumulative division/replacement counts **(J)**, allele counts **(K)**, percentage of singletons among all alleles **(L)**, allele birth counts **(M)** and allele death counts **(N)**. Allele death counts (lines) are categorized into mutation events (circles) and division/replacement events (diamonds). Dark blue = Moran A, green = Moran B, yellow = “binomial BP” with *g*_0_ = *g*_2_ = 0.25 (non-extinction), orange = “binomial BP” with *g*_0_ = *g*_2_ = 0.25 (*N* (*t*_*f*_) *∈* [90, 110]), purple = “fast BP” with *g*_0_ = *g*_2_ = 0.5 (*N* (*t*_*f*_) *∈* [90, 110]), cyan = “slow BP” with *g*_0_ = *g*_2_ = 0.05 (*N* (*t*_*f*_) *∈* [90, 110]), gray = “supercritical BP” with *g*_0_ = 0.2465, *g*_1_ = 0.5 and *g*_2_ = 0.2535 (non-extinction).

Because condition (2) is no longer satisfied, the fitness coefficients and mutation rates are not at equilibrium and fitness in Moran A increases over time (**Figure 5I**), leading to an increase in its replacement count (**Figure 5J**) as compared to the neutral or balanced evolution setting. As a result, the counts of both alleles and singletons are slightly lower in the selective evolution as compared to previous settings (**Figure 5B-C**). The same is also true for all remaining models. Remarkably, binomial BP still behaves identically to Moran A, differing only in population size over time (**Figure 5G**). As before, relaxing the conditioning on binomial BP leads only to higher variances of the statistics without changing their averages. Like in previous examples, supercritical BP has much higher averages and variances in all statistics than other models, both at the end of simulation as well as throughout time. Similarly as in the previous case, changing the progeny cell count distribution in BP while retaining criticality results in allele count and singleton count converging to different values (**Figure 5B-C, K-L**).

As in the balanced evolution setting, the only model differing in fitness from the remaining critical processes is Moran B (**Figure 5I**), this time resulting in twice as many replacements compared to other models (**Figure 5F, J**). In later moments of the simulation, the number of replacements is even higher than in the critical BP. Consequently, in Moran B, the counts of alleles and singletons decrease at a fast rate after reaching maximum values (**Figure 5B-C, K-L**).

#### 3.1.4 Passenger domination case

Finally, we investigate the setting where passenger mutations are strongly deleterious, with parameters *s* = 0, *d* = 0.5, *μ*_*d*_ = 0.1 and *p* = 1*/*10 corresponding to **Figure 6**.

**Figure 6:**
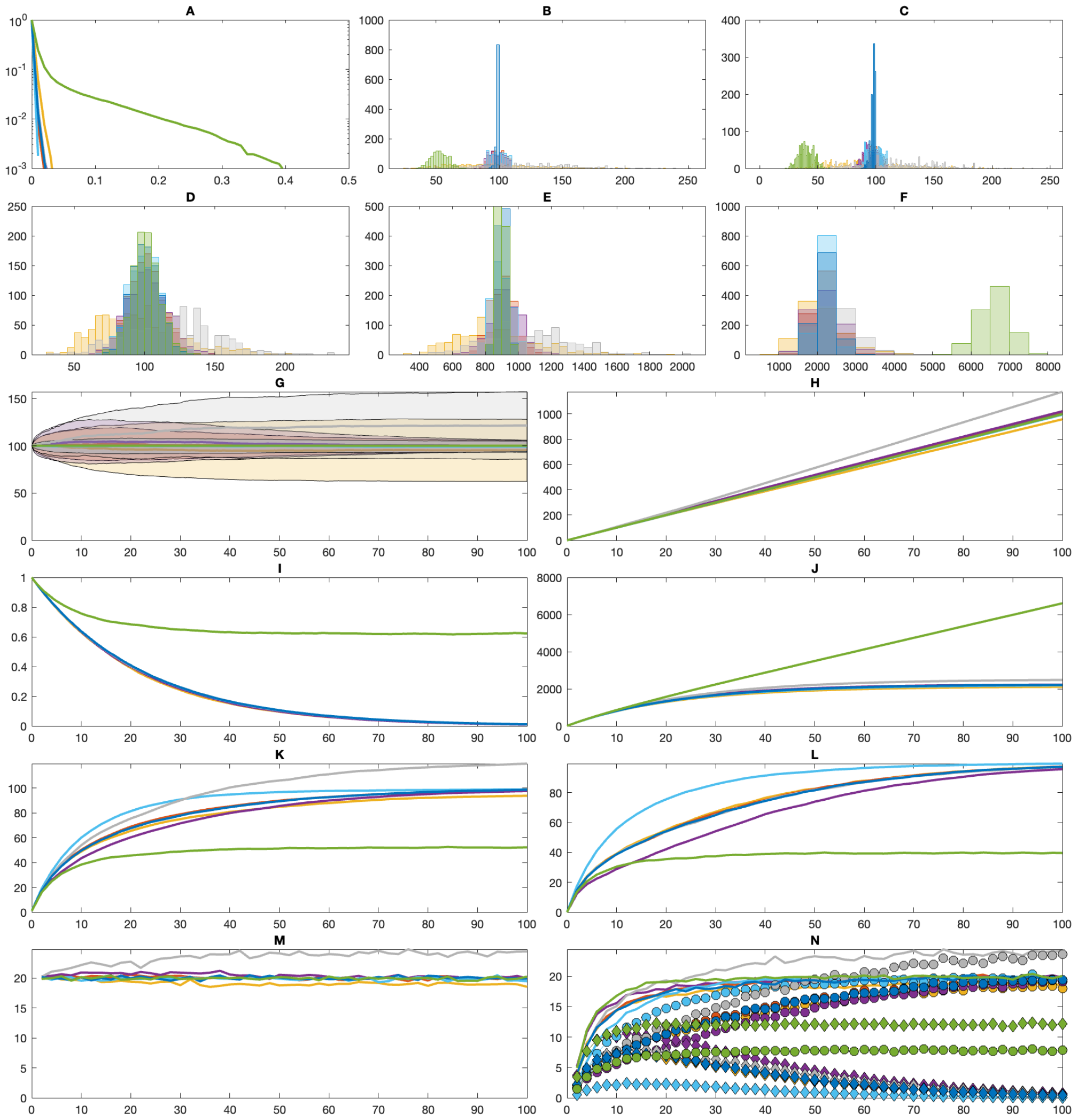
Comparisons between Moran and branching process (BP) models in the “deleterious” setting. **(A)** Average cumulative tail of the mutational Site Frequency Spectra. **(B)** Distributions of allele counts at *t*_*f*_. **(C)** Distributions of singleton counts at *t*_*f*_. **(D-F)** Distributions of counts of driver mutations **(D)**, passenger mutations **(E)** and divisions **(F)** within [*t*_0_, *t*_*f*_]. **(G-N)** Trajectories of the averages over time of population sizes (+/-std) **(G)**, cumulative mutation counts **(H)**, fitness **(I)**, cumulative division/replacement counts **(J)**, allele counts **(K)**, percentage of singletons among all alleles **(L)**, allele birth counts **(M)** and allele death counts **(N)**. Allele death counts (lines) are categorized into mutation events (circles) and division/replacement events (diamonds). Dark blue = Moran A, green = Moran B, yellow = “binomial BP” with *g*_0_ = *g*_2_ = 0.25 (non-extinction), orange = “binomial BP” with *g*_0_ = *g*_2_ = 0.25 (*N* (*t*_*f*_) *∈* [90, 110]), purple = “fast BP” with *g*_0_ = *g*_2_ = 0.5 (*N* (*t*_*f*_) *∈* [90, 110]), cyan = “slow BP” with *g*_0_ = *g*_2_ = 0.05 (*N* (*t*_*f*_) *∈* [90, 110]), gray = “supercritical BP” with *g*_0_ = 0.2465, *g*_1_ = 0.5 and *g*_2_ = 0.2535 (non-extinction).

In Moran A, as cells accumulate increasingly more mutations, their fitness decrease to 0 because of the passenger mutations’ deleterious coefficient (**Figure 6I**), therefore they stop dividing (**Figure 6J**). However, the mutation process depends only on the population size (**Figure 6G**) and therefore occurs at a constant rate throughout time (**Figure 6H**). The consequence is that every cell sooner or later would acquire a unique mutation, therefore the cell population almost consists only of singletons (**Figure 6B-C**).

As is the case for other settings, binomial BP matches the average statistics from Moran A throughout history and at the final time (Appendix B, Table 5). The same is true for relaxing the conditioning on binomial BP, which only increases the variances. However, unlike the selective evolution setting, altering the progeny cell count distribution does not change the steady state values for the allele and singleton counts, as both converge to the same distributions as binomial BP and Moran A (**Figure 6B-C, K-L**). Nonetheless, compared to these models, the fast BP converges faster and the slow BP takes more time to converge.

**Table 5:** Statistics for Figure 6 (passenger domination)

| Model | B | C | D | E | F |
| --- | --- | --- | --- | --- | --- |
| Dark blue | 98.5±1.2 | 97.3±2.3 | 99.7±10.0 | 901.1±28.7 | 2216.4±236.6 |
| Green | 52.2±6.6 | 39.5±6.4 | 100.5±9.8 | 898.1±28.8 | 6610.3±395.6 |
| Yellow | 93.7±32.0 | 92.4±31.4 | 96.3±29.2 | 862.2±241.9 | 2106.2±579.5 |
| Orange | 98.4±5.9 | 97.1±6.1 | 100.0±12.1 | 904.2±75.6 | 2187.1±301.4 |
| Purple | 97.5±6.1 | 95.5±6.5 | 101.7±14.2 | 920.7±92.6 | 2226.7±406.0 |
| Cyan | 98.8±5.9 | 98.5±6.0 | 99.1±11.1 | 895.5±58.5 | 2183.8±186.4 |
| Gray | 119.6±34.7 | 117.9±34.1 | 117.3±29.4 | 1059.3±255.4 | 2477.0±573.8 |

The deleterious evolution setting is the only scenario in which supercritical BP behaves similarly to most of the other models. While fitness tends to zero, cells stop dividing an the impact of the supercriticality is no longer significant.

Finally, as usual, Moran B has a much higher steady state fitness compared to other models (**Figure 6I**). Therefore, although the replacement count is lower than in the balanced or selective evolution settings, cells do not stop dividing as is the case with Moran A, because the fitness does not converge to 0 (**Figure 6E, I**). This results in much lower allele count and singleton count (**Figure 6A-B, J-K**). Despite accumulating passenger mutation, after the initial period of dropping the average fitness stays at nonzero value, due to the drift process favoring fixation of clones with higher fitness.

In the deleterious evolution scenario, since cells stop dividing, the main reason of cells’ death is mutation (**Figure 6N**), except model B in which cells keep dividing (**Figure 6J**).

### 3.2 Fitting breast cancer SFS

We use the Moran A model to fit the mutational SFS from 3 samples of breast cancer. We fix population size *N* = 100 cells, average time between mutation events *L* = *Nμ* = 6, probability of driver mutations *p* = 0.01, final time *t*_*f*_ = 100. We then vary the values for *s* and *d*, simulate the SFS from 1,000 simulations and compute the average SFS. For a given sample, we compute the cumulative tail of the SFS *S*(*f*_*i*_) i.e., the proportion of mutations occurring at frequencies *> f*_*i*_, and similarly the average {*S*(*f*_*i*_|*s, d*)} for every combination of (*s, d*). The reverse cumulative SFS is evaluated for mutations with frequency *f >* 0.05. The error for (*s, d*) is defined as

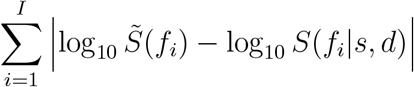

where *I* is the largest index such that 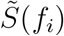 and *S*(*f*_*i*_|*s, d*) are both positive.

Figures 7-9 present the fitting results for the SFS from the breast SFS data. The (*s, d*) combinations with low error exhibit a trade-off between driver and passenger mutations: the observed SFS can be simulated by Moran A with either low values for *s* and *d*, or high values for both (panel A in each figure). As a result, the 100 best (*s, d*) parameters (marked as squares) can be approximated by linear regression.

**Figure 7:**
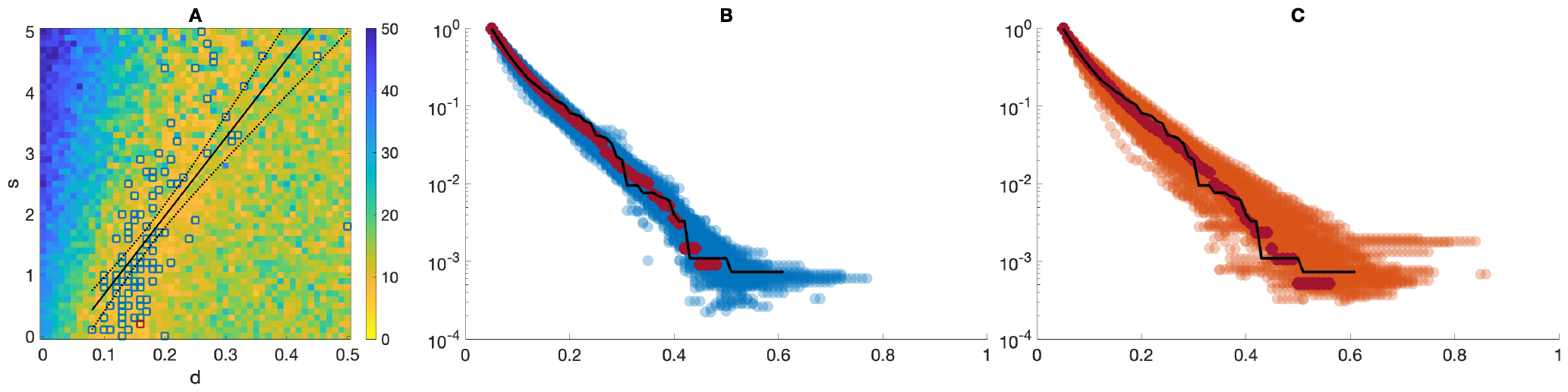
Results from fitting the SFS from sample G2. **(A)** Heatmap of error between data SFS tail and average SFS tail for distinct (*s, d*) combinations in Moran A. Dark blue squares: 100 best (*s, d*) combinations, with linear regression. Dark red square: best (*s, d*) combination. **(B)** Comparison between sample SFS (black line), average SFS under Moran A from 100 best (*s, d*) combinations (dark blue dots) and the best (*s, d*) combination (dark red dots). **(C)** Comparison between sample SFS (black line), average SFS under “binomial BP” (*g*_0_ = *g*_2_ = 0.25, *N* (*t*_*f*_) *∈* [90, 110]) from 100 best (*s, d*) combinations (red dots) and the best (*s, d*) combination (dark red dots). Each SFS from Moran A or binomial BP is averaged from 1,000 simulations.

The range of best *d* parameter values does not vary among cases, with the G2 sample (Figure 7, panel A) having greater tolerance for changes in this parameter value. The impact of the *s* parameter on the SFS tail shape is significant: the range of the best fits varies between cases. The value of *s* is small for sample G2 (Figure 7, panel A) and slightly bigger for G32 (Figure 8, panel A). In the case of sample G41, the best fit was obtained using a relatively high *s* parameter value (Figure 9, panel A), indicating strong selection.

**Figure 8:**
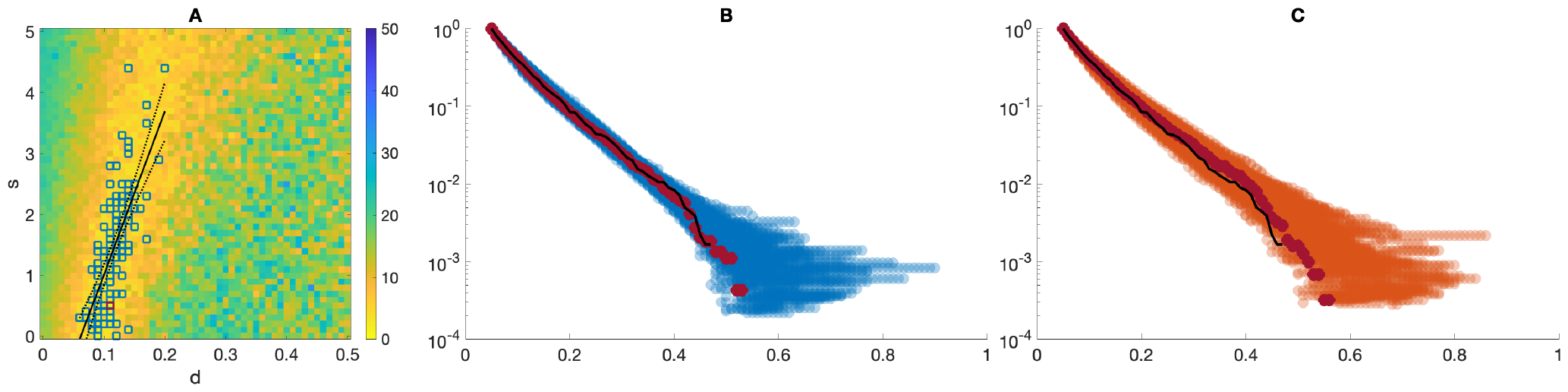
Results from fitting the SFS from sample G32. **(A)** Heatmap of error between data SFS and average SFS for distinct (*s, d*) combinations in Moran A. Dark blue squares: 100 best (*s, d*) combinations, with linear regression. Dark red square: best (*s, d*) combination. **(B)** Comparison between sample SFS (black line), average SFS under Moran A from 100 best (*s, d*) combinations (dark blue dots) and the best (*s, d*) combination (dark red dots). **(C)** Comparison between sample SFS (black line), average SFS under “binomial BP” (*g*_0_ = *g*_2_ = 0.25, *N* (*t*_*f*_) *∈* [90, 110]) from 100 best (*s, d*) combinations (red dots) and the best (*s, d*) combination (dark red dots). Each SFS from Moran A or binomial BP is averaged from 1,000 simulations.

**Figure 9:**
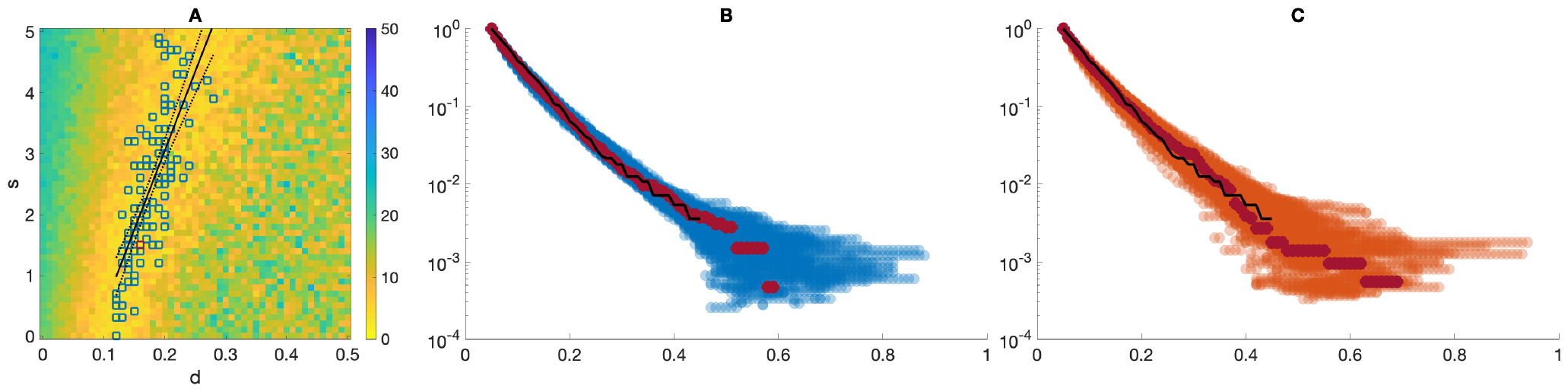
Results from fitting the SFS from sample G41. **(A)** Heatmap of error between data SFS and average SFS for distinct (*s, d*) combinations in Moran A. Dark blue squares: 100 best (*s, d*) combinations, with linear regression. Dark red square: best (*s, d*) combination. **(B)** Comparison between sample SFS (black line), average SFS under Moran A from 100 best (*s, d*) combinations (dark blue dots) and the best (*s, d*) combination (dark red dots). **(C)** Comparison between sample SFS (black line), average SFS under “binomial BP” (*g*_0_ = *g*_2_ = 0.25, *N* (*t*_*f*_) *∈* [90, 110]) from 100 best (*s, d*) combinations (red dots) and the best (*s, d*) combination (dark red dots). Each SFS from Moran A or binomial BP is averaged from 1,000 simulations.

We compare the SFS from the 100 best (*s, d*) parameters against the breast cancer data-based SFS (Figures 7-9, panel B). The SFS from each sample is well fitted with Moran A, not only with the optimal parameter set but also with other (*s, d*) combinations. The fit is particularly good for the region of SFS with low frequency (*f <* 0.2). There are exponentially fewer mutations occurring at larger *f*, resulting in relatively higher discrepancy in the long tail of the SFS from Moran A. However, the overall shape of the observed SFS can be fitted well by the Moran A model.

We also compare the SFS from the binomial BP (*g*_0_ = *g*_2_ = 0.25, *N* (*t*_*f*_) *∈* [90, 110]), using the 100 best (*s, d*) parameters from Moran A inference (Figures 7-9, panel C). Similarly to Moran A, the SFS from BP can fit the shape of the observed SFS tail well. This is consistent with our finding from the previous section that the binomial BP with tight conditioning on the final population size behaves similarly to the Moran A model, resulting in similar statistics under a range of selection scenarios.

## 4 Discussion and Conclusion

As it has been expected, Moran model A behaves comparably to the binomial BP conditioned on final population size being close to the initial count. This manifests in similar statistics under different extremes of selection scenarios (Section 3.1), including values that are observable in sequencing samples. Crucially, SFS fitting results for experimental breast cancer data (Section 3.2) are similar between the two models. This finding might hold mathematical importance, and requires further investigation. Moreover, interestingly, the similarity between Moran A and binomial BP becomes more pronounced as the latter is conditioned more tightly to resemble the constant population size expected in Moran models. When conditioned only on non-extinction, the population size of each BP realization may deviate significantly (**Figures 3-6G**). This leads to higher variances in allele and singleton counts (**Figures 3-6B-C, K-L**) as well as mutation and division/replacement counts (**Figures 3-5D-F, H-N**), although the means of these statistics remain similar (Appendix B, Tables 2-5). However, the high population size variance also results in different SFS in non-extinction BP compared to tightly conditioned BP and Moran model A (**Figures 3-6A**).

Our Moran model B, similar though not identical to the model introduced in [2], exhibits a phenomenon known as the drift barrier, which prevents the deleterious passenger mutations from dominating temporal trends in fitness, even under mutation-selection balance condition tipped in their favor (*sp < dq*). Indeed, under this condition, due to drift, the fitness may decrease or increase depending on how much smaller *sp* is than *dq*. In addition, fitness generally increases at mutation-selection equilibrium (*sp* = *dq*). On the contrary, trends in Moran model A and binomial BP model follow the mutation-selection condition.

The effect of increasing fitness in model B was already predicted in [2] and described in [16]. While in Moran A fitness stays constant (as expected), in the case of Moran B, clones with higher fitness are favored, even for the same initial conditions and in absence of new mutations (see section 3.1.1 in [16]). This behavior results from the difference between Moran A and Moran B in the expected change in population fitness after a death-replacement event. As shown in Eqs. (4) and (5) in [16], the expected fitness change is 0 in Moran A and is *≥* 0 in Moran B. In general, fitness in Moran A depends only on the balance between drivers and passengers, while the trends in Moran B are more complex, as explained mathematically and confirmed by simulations in [2]. The drift and selection pattern in Moran B biases it toward increasing fitness.

Among the BP variations, the supercritical model behaves differently in all cases compared to other models, due to the difference in population size growth rate (**Figures 3-5G**). The only exception is the deleterious evolution setting, in which the impact of supercriticality is less prominent, since the fitness being close to zero means cells stop dividing. In this scenario all statistics are much more similar to those obtained from other models (Appendix C, Table 5).

Our experiments with different progeny cell count distributions in BP show that the fast BP always has higher variances in the population size throughout time compared to the binomial BP, even if similarly conditioned (**Figures 3-6G**). The fast BP also results in both less alleles and lower percentage of singletons within all alleles (**Figures 3-5B-C, K-L**). This is not observed in case of deleterious evolution (**Figure 6B-C, K-L**), where there is only a difference in the rate of reaching the steady state, which is the same as for other models. On the other hand, the averages of mutation and division/replacement event counts (**Figures 3-6D-F** and Appendix C, Table 2-5) do not differ from averages for Moran model A and binomial BP. Reversely, the population size in slow BP varies less, and the allele count and percentage of singleton count is much higher, compared to the binomial BP.

There are features shared between all Moran models and BP variations across different selection scenarios. The more division/replacement events occur during a simulation, the less alleles and singletons we observe both at the sampling time point as well as throughout tumor history. This is especially pronounced in case of deleterious evolution, where in Moran B continuously dividing population under selective pressure prevents the accumulation of singletons (**Figure 6B-C, K-L**). Conversely, if the events occurring during a simulation are dominantly mutations, then the population consists of more alleles and singletons. In conclusion, across models, higher selection is associated with less alleles and singletons, higher pace of allele death, and cumulative SFS with fat tail.

It seems relevant to note that the frequently cited reference by Gerrish and Lenski [11] introduces a model of competition in populations of constant size in an asexual population. From the reading of this corner-stone paper, it seems that it uses results from supercritical branching processes and then just scales them intuitively into the constant population size framework. This method seems not mathematically rigorous. Our comparison of Moran and branching process models identifies subtle but important differences between the two approaches. Overall, we have shown that the critical binomial BP and the Moran A model behave similarly in the Tug-of-War setting under distinct selection scenarios. This finding is relevant for improving simulating efficiency and optimizing model inference. Branching process and Moran model remain the two main stochastic modeling approaches in population genetics, where they provide the theoretical framework to uncover a tumor’s history from sequencing snapshots. However, BP simulation is considerably more time-consuming, as the cell population size can change arbitrarily due to random fluctuations. This problem is exacerbated in critical or near-critical BP, which is applicable for modeling many cancers. In this setting, the BP often has high probability of extinction, hence the high fraction of simulations that have to be discarded makes model inference computationally costly. In such cases, it would be more efficient to employ Moran A, which we have shown to provide comparable sample statistics and which is easier to implement. However, more work is needed to establish the theoretical equivalence between the Moran A model and the critical BP, and if this compatibility breaks down under certain conditions.

Both the Moran A model and the binomial BP can fit the SFS tail in our breast cancer samples well. However, the inference is complicated by a wide range of selection coefficients that result in equally comparable SFS to the data. These coefficients exhibit a trade-off between driver and passenger mutations, as the same SFS can result from driver mutations being more advantageous if the passenger mutations are also more deleterious, and vice versa. Therefore, the mutational SFS alone is not adequate to differentiate between these different selection settings. Separately, we found that the inference for all of our samples requires *d >* 0, confirming the observations from McFarland et al. [19] that passenger mutations exhibit a deleterious effect during tumor progression.

## Data availability

The breast cancer sequencing data can be found under https://ega-archive.org/ with accession number: EGAD00001009081. Any queries should be directed to the corresponding author.

## APPENDIX

### A Elements of mathematical population genetics

#### A.1 Wright-Fisher model and Moran model comparison

##### A1.1 Wright–Fisher model

There are two copies of each gene in a diploid cell. The copies can be of the same allele (e.g. *AA* or *aa*) or two different alleles (*Aa*). If the population consists of *N* diploid individuals, there exist 2*N* copies in total. In the accordance with Wright-Fisher model, in each generation random alleles are drawn with replacement from gene pool of size 2*N* – the generations do not overlap. Such reproduction scheme can be described mathematically as a discrete-time Markov chain – the future allele frequencies are dependent only on the present frequencies, not on those from past generations. Due to the randomness in the process, allele frequencies change at a rate which is inversely proportional to the population size. These fluctuations correspond to a process of genetic drift.

The state of the population in each generation can be described as the number of *A* alleles in the population, which can range from 0 (loss of allele *A*) to 2*N* (fixation of allele *A* and loss of allele *a*). The states 0 and 2*N* are called “absorbing states” because the population is not able to leave any of these (considering no mutation or migration events). In other cases, the transition probability can be calculated based on binomial distribution. In the population with *i* copies of allele *A*, the frequency of allele *A* is equal to *p* = *i/*2*N* and the frequency of allele *a* is *q* = 1 *− p*. The probability of changing state from *i* copies of *A* to *j* copies of *A* (for *i, j* = 0, 1, 2, …, 2*N*) in one generation is [13]:

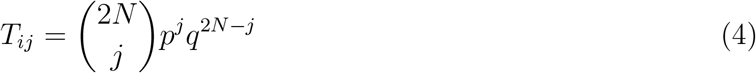

#### Moran model

In the Moran model [21], in each step of algorithm, a single random allele *x* from haploid population of 2*N* individuals dies and is being “replaced” by another randomly chosen allele from the population (including *x* itself), thus ensuring that the population size remains constant. Unlike the Wright-Fisher model, where an allele can have up to 2*N* offspring, in the Moran model the allele can have 0 or 2 descendants. The time between birth-death events (the lifespan of an individual allele) is exponentially distributed with mean equal to 1 and generations do overlap.

As in the Wright-Fisher model, this reproduction scheme can be described mathematically by Markov chain. In this case, however, it is the continuous-time Markov chain with values in the set {0, 1, …, 2*N }*. Therefore, the mathematical solutions for the Moran model are easier to ascertain than the Wright-Fisher model. However, there are fewer time steps to be computed in the WrightFisher model. As a result, the simulations are more computationally convenient in the Wright-Fisher framework, compared to the Moran model. The Moran and Wright-Fisher models give qualitatively similar results, but genetic drift runs twice as fast in the Moran model [9].

The details regarding Moran model are described in section 2.1.

#### A.2 Infinitely many alleles version of Wright-Fisher model

For neutrality testing purposes we consider the “infinitely many alleles” (IAM) version of the Wright-Fisher model. This type of model is particularly useful in molecular population genetics and was inspired by molecular nature of the gene. Average gene is sequence of 3000 nucleotides (A, G, T and C), so there are 4^3000^ possible sequences (alleles), the number which for practical purposes can be taken as infinity, what leads to infinitely many alleles model.

Most nucleotide mutations will lead to sequences not currently existing in the population, so in this case all mutants are assumed to be of a new allelic type - there is no reverse mutation. In such a model each allele will sooner or later be lost from the population.

#### A.2.1 Expected allele number

Under assumptions of the IAM, let us define **A** = (*A*_1_, *A*_2_, …, *A*_*n*_) as the vector of the allelic types each of which is represented by exactly *j* genes in the sample. The following Ewens Sampling Formula (ESF) was derived in [10] and [14]

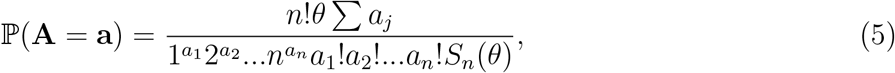

with

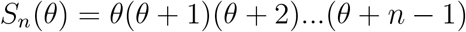

Furthermore, the probability distribution of number *K* of different alleles in the sample has the form

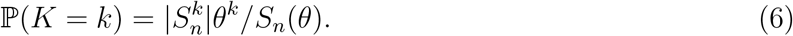

Under neutrality *θ* = *nμ*, where *μ* denotes mutation coefficient and *n* is the sample size. 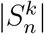is the absolute value of a Stirling number of the first kind. The expression for the expected value of *K* can be derived from Equ. (6)

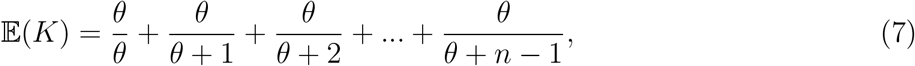

with the corresponding expression for variance of *K*

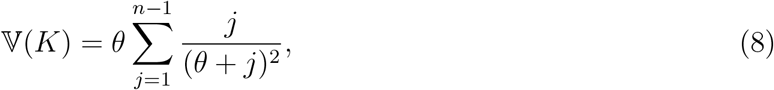

Equations (5) and (6) show jointly that the conditional distribution of the vector **A** = (*A*_1_, *A*_2_, …*A*_*n*_), given the value of *K*, is

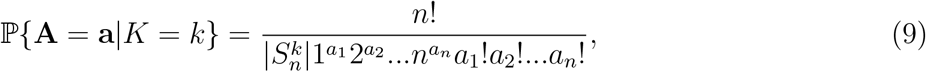

where **a** = (*a*_1_, *a*_2_, …, *a*_*n*_). From (9) the procedure can be derived, which allows testing the null hypothesis that the alleles in the sample are selectively equivalent (neutral).

### B Statistics in Section 3.1

Tables 2, 3, 4, 5 contain some statistics for Figures. 3, 4, 5, 6, respectively. The models are named according to their colors in the figures, which are listed in legend for Table 2 along with the statistics.

